# Efficient curation of genebanks using next generation sequencing reveals substantial duplication of germplasm accessions

**DOI:** 10.1101/410779

**Authors:** Narinder Singh, Shuangye Wu, W. John Raupp, Sunish Sehgal, Sanu Arora, Vijay Tiwari, Prashant Vikram, Sukhwinder Singh, Parveen Chhuneja, Bikram S. Gill, Jesse Poland

## Abstract

Genebanks are valuable resources for crop improvement through the acquisition, *ex-situ* conservation and sharing of unique germplasm among plant breeders and geneticists. With over seven million existing accessions and increasing storage demands and costs, genebanks need efficient characterization and curation to make them more accessible and usable and to reduce operating costs, so that the crop improvement community can most effectively leverage this vast resource of untapped novel genetic diversity. However, the sharing and inconsistent documentation of germplasm often results in unintentionally duplicated collections with poor characterization and many identical accessions that can be hard or impossible to identify without passport information and unmatched accession identifiers. Here we demonstrate the use of genotypic information from these accessions using a cost-effective next generation sequencing platform to find and remove duplications. We identify and characterize over 50% duplicated accessions both within and across genebank collections of *Aegilops tauschii*, an important wild relative of wheat and source of genetic diversity for wheat improvement. We present a pipeline to identify and remove identical accessions within and among genebanks and curate globally unique accessions. We also show how this approach can also be applied to future collection efforts to avoid the accumulation of identical material. When coordinated across global genebanks, this approach will ultimately allow for cost effective and efficient management of germplasm and better stewarding of these valuable resources.

## INTRODUCTION

With an estimate of more than 1 billion underfed people in the world and projected human population growth to over 9 billion by 2050^1^, there is increased food insecurity risk and an even a greater challenge to global food supply. To meet the future demand food production needs to be doubled^2,3^ in the midst of shrinking resources^4^. A critical raw ingredient for continued crop improvement is genetic diversity. Although flowering plants have a huge diversity, mankind cultivates only a handful of them for food and feed with about 90% of the food and feed coming from only ten cultivated crop species^5,6^. Great opportunities exist to domesticate new plant species and improve the existing crop plants^7^. Genetic diversity present in wild crop relatives and that conserved in genebanks are a source of novel genes that increase yield, resistance to pests and disease and abiotic stress.

Genebanks play an imperative role in *ex-situ* germplasm conservation that is critical for crop improvement. These facilities provide infrastructure for storage, a platform for sharing, and opportunity for better access and utilization of the germplasm. More than 1700 genebanks around the world stock over 7 million plant accessions^8^, of which only a small number are characterized and few are ever used for crop improvement^9^. Although genebanks are crucial for aforementioned reasons, they are expensive to establish and manage^9^. Therefore, to maximize the value of this investment and of the germplasm resources, strategies for efficient genebank management are needed.

Researchers have implemented different strategies to prioritize a limited number of potentially useful accessions from genebanks that can be used for crop improvement. These strategies include selecting accessions based on their phenotype and associated passport data. One example of such strategies is Focused Identification of Germplasm Strategy (FIGS) that works on the premise that the adaptive traits shown by the accessions is the direct result of environmental conditions of their respective place of origin, and the genetic diversity can be maximized by sampling accessions based on their diverse contrasting geographic regions^10,11^. However, accessions stored in the genebanks are often missing the phenotypic and passport data, or could be associated with incorrect passport data, which limits the application of FIGS. Other limitations of such strategies include the high cost of phenotyping, limited resources such as space and manpower to do such screening on a larger scale. Therefore, cheaper and reliable methods that are free from these kinds of uncertainties are needed.

Contrary to the unreliable phenotypic and passport information, genotypic characterization of accessions should provide better curation of genebanks and optimize the use of genetic diversity. Modern tools and techniques such as next-generation sequencing (NGS) and genotyping-by-sequencing (GBS) can be used to rapidly and cost-effectively characterize germplasm stored in genebanks^12^. Data generated by this approach can be used for identifying identical accessions (duplications) within and among genebanks, characterizing genomic diversity^13^, and imputing missing passport information. Identifying and removing identical accessions from genebanks reduces the cost while increasing the efficiency of managing and utilizing genebank resources.

Consortiums such as the DivSeek initiative (http://www.divseek.org) exist with a vested interest in genotyping the germplasm stored in genebanks for the purpose of genetically characterizing these resources and optimizing the use of the genetic diversity. The Wheat Genetics Resource Center (WGRC; http://www.k-state.edu/wheat-iucrc), an NSF Industry/University Cooperative Research Center, located at Kansas State University in Manhattan, KS, USA, is another example of such effort to characterize wild species stored in the in-house and collaborative genebanks. WGRC primarily specializes as a working collection of wheat genetic diversity and focuses on collecting, evaluating, identifying and mobilizing the genetic diversity. Other major genebanks are managed by the Consultative Group on International Agriculture Research (CGIAR) center throughout the world such as the International Maize and Wheat Improvement Center (CIMMYT; Mexico). CIMMYT holds over 105,000 Triticeae accessions in their global genebank outside of Mexico City. Another important CGIAR genebank, International Center for Agriculture Research in Dry Areas (ICARDA), with over 41,000 Triticeae accessions, was recently relocated to Terbol, Lebanon and Rabat, Morocco due to inaccessibility of original collection in Aleppo, Syria^14^ (www.genebanks.org). Systematic safety backup at Svalbard Global Seed Vault ensured the timely restoration of germplasm at ICARDA. Several other genebanks at regional and national level throughout the world, such Punjab Agricultural University (PAU; Ludhiana, India), carry accessions of local importance that are utilized for germplasm improvement and breeding. With these genebanks operating individually and in conjunction with each other, it becomes imperative to understand the status of shared and duplicated accessions within and across these genebanks.

Modern hexaploid bread wheat (*Triticum aestivum* L.) is a critical focus to mitigate the upcoming food security challenge in coming decades. In the context of continued wheat improvement through breeding, maintaining and increasing genetic diversity in wheat is very important. Due to genetic bottlenecks from domestication and modern breeding, wheat has a limited genetic base. Its domestication coexisted with the advent of agriculture about 10,000 years ago^15-18^. Three distinct diploid species—*Triticum urartu* (AA), a relative the extant *Aegilops speltoides* (BB), and *Aegilops tauschii* (DD)—contributed to the origin and evolution of polyploid wheat (AABBDD). First natural hybridization of *Triticum urartu* and B-genome donor resulted in tetraploid *Triticum turgidum* (AABB) wheat around 0.58-0.82 million years ago^19^ followed by a second whole-genome hybridization with *Aegilops tauschii* (DD)^20,21^ in the fertile crescent around the Caspian sea, to give rise to modern hexaploid wheat. The limited hybridization with *Ae. tauschii*, followed by domestication and improvement has severely limited the genetic diversity of the wheat D genome^22^. The presence of great genetic diversity in these wild relatives provides an excellent resource for continued improvement.

As a proof of concept for genebank curation, we used *Ae. tauschii* as a model for this study while providing valuable and needed curation of several important repositories for this species. The main objectives of this study were to (i) genotype the entire collections of *Ae. tauschii* from three different genebanks using a cost effective and robust reduced representation sequencing, (ii) identify identical accessions within genebanks using genotypic data, (iii) identify identical accessions between genebanks using genotypic data, and (iv) develop protocols for efficiently curating genebanks.

## METHODS

### Germplasm acquisition

A total of 1143 accessions of *Ae. tauschii* were assessed, which included 568 accessions from the Wheat Genetics Resource Center (WGRC, Kansas State University), 187 accessions from Punjab Agricultural University (PAU; Ludhiana, India), and 388 accessions from Centro Internacional de Mejoramiento de Maíz y Trigo (CIMMYT; Mexico) (Supplementary Table S1). The germplasm consisted of the accessions collected from natural habitat (Fig. 1) and accessions received from other genebanks.

**Figure 1.**
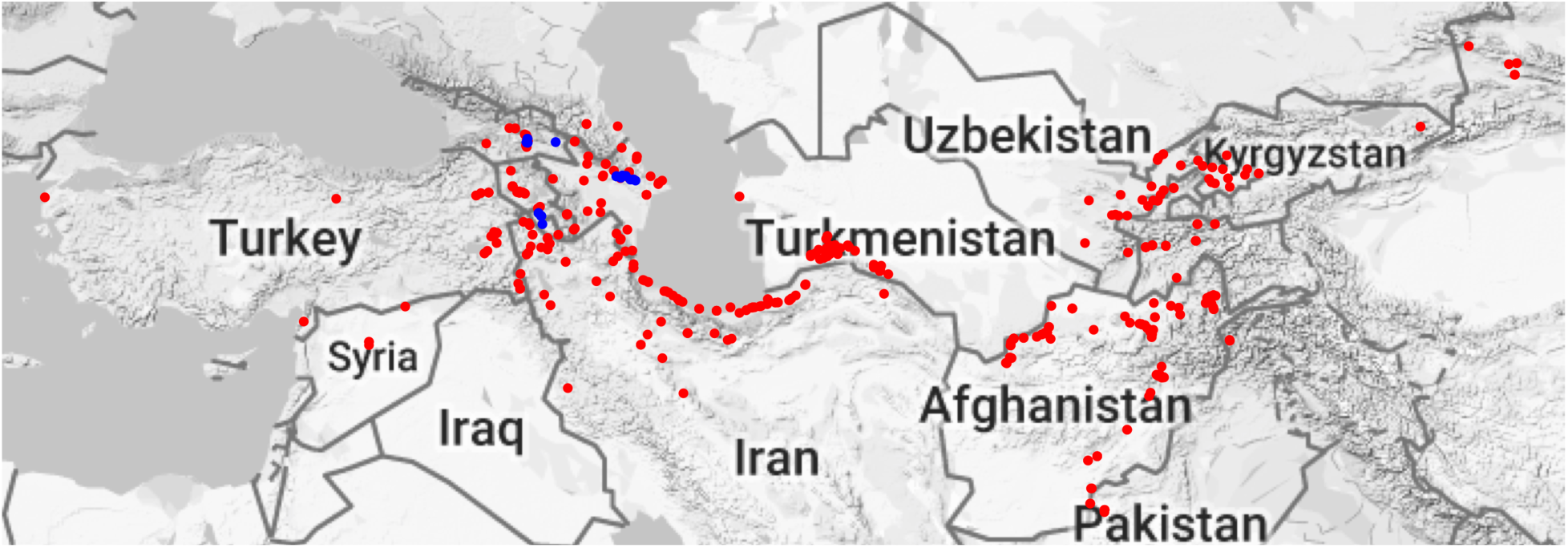
Geographical distribution of the WGRC accessions. Each dot represents a collection site for *Aegilops tauschii* accessions. Blue dots represent newly collected accessions (June 2012), and red dots represents previously collected accessions (1950s and 60s). Two accessions from China’s Shaanxi and one from Henan are not shown here to control for the size of the map.

### DNA extraction and Genotyping

Two approaches for DNA extraction and the GBS libraries preparation were implemented for WGRC and PAU accessions (hereafter referred to as Set 1), and CIMMYT accessions (hereafter referred to as Set 2). For Set 1, young leaf tissues from 2-3 weeks old seedlings were collected in 96 well plates. Genomic DNA was extracted using Qiagen BioSprint 96 DNA Plant Kit (QIAGEN, Hilden, Germany) and quantified with Quant-iT(tm) PicoGreen^®^ dsDNA Assay Kit (ThermoFisher Scientific, Waltham, MA, USA). At least one random well per plate was left blank with known position for quality control and library integrity. GBS libraries were prepared following the protocol from Poland *et al*. (2012). Briefly, the libraries were prepared in 95-plex using 384A adapter set. For complexity reduction, DNA for each sample was digested using two enzymes – rare cutter *PstI* (CTGCAG), to which the uniquely barcoded adaptors were ligated, and frequent cutter *MspI* (CCGG), to which the common reverse adapter was ligated. All samples from a single plate were pooled and amplified using polymerase chain reaction (PCR). Detailed protocol can be found at http://wheatgenetics.org/download/category/3-protocols. Libraries were sequenced on ten lanes in total on Illumina HiSeq2000 (Illumina, San Diego, CA, USA) platform at University of Missouri (UMC; Columbia, Missouri) and McGill Univesity-Génome Quebec Innovation Centre (Montreal, Canada) facility. To compute the error rate for the GBS, 76 WGRC accessions were randomly chosen, and were sequenced as biological replications (different seedlings) using the abovementioned protocol.

For Set 2, *Ae. tauschii* accessions were planted in greenhouse in plots. Leaves from single seedling plants were taken and DNA was extracted using modified CTAB method^24^ and quantified using NanoDrop spectrophotometer V2.1.0 (ThermoFisher Scientific, Waltham, MA, USA). Genotyping was performed at DArT, Canberra, Australia using DArTseq^25^ methodology that has been used in recent years at CIMMYT^25-27^. DArTseq is a combination of diversity array technology (DArT)^28,29^ complexity reduction and next-generation sequencing (NGS) methods. Two optimized enzyme sets, *PstI*-*HpaII* and *PstI*-*HhaI*, were used for complexity reduction. Samples were sequenced twice using two different 4bp cutters on one end of the RE fragments (*HpaII* and *HhaI*) on a total of nine lanes.

### Single Nucleotide Polymorphism discovery

Single nucleotide polymorphisms (SNPs) were discovered and typed with TASSEL-GBS^30^ framework (http://www.maizegenetics.net) using an in-house written Java plugin and a modified Java pipeline without reference genome. In brief, 64bp long valid tags (containing restriction cut site and a barcode) were extracted from each sample, and then similar tags (up to 3bp differences) were internally aligned to find SNPs. To test putative tag pairs for as allelic SNP calls, Fisher exact test was performed on all aligned tag pairs with one to three nucleotide differences. Tag pairs that failed the test at *P* ≤ 0.001 were considered biallelic and converted to SNP calls^31^. As the accessions are inbred lines, this test determined allelic tags that are disassociated (e.g. only one of the two alternate tags present in any given individual) and can be considered alternate tags for SNP alleles at the same locus. Due to the differences in library preparation for Set 1 and Set 2, the tag discovery step was performed using only Set 1 accessions, and then the discovered tags were used as reference to produce SNPs for both sets.

### Statistical analyses, allele matching and error computation

Data analyses and genotype curation were performed using custom scripts in *R* statistical language^32^ to find identical accessions within and among genebanks. In addition to hierarchical clustering (Supplementary Fig. S1), an identity matrix was computed by pairwise comparison of accessions across all SNP sites. Hierarchical clustering group individuals based on the relative genetic distance between individuals, whereas, pairwise allele matching provides an absolute percent identity by state (IBS) coefficient between all individuals. Although, clustering can provide an independent support for allele matching, it is hard to interpret clustering to identify identical accessions. However, clustering can provide a quick method to identify obvious outliers and misclassified accessions (Supplementary Fig. S1). For clustering, population-level SNP filtering was performed to retain the SNPs with ≤50% missing data. In contrast, for pairwise comparison, only those SNP sites without missing data and homozygous in both individuals were used for comparison. A stringent threshold of 99% identity was used to consider two accessions the same to account for a 1% sequencing and alignment error rate. Accessions with ≥99% identity were considered identical within and/or across genebanks. Percent Identity by State (pIBS) was computed using the following equation 1:

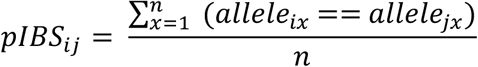

where, *pIBS*_*ij*_ is the percent Identity by State for a given pair of accessions *i* and *j*, *allele*_*ix*_ and *allele*_*jx*_ are the alleles at *x*^*th*^ SNP of accessions *i* and *j*, respectively, == sign represents an exact successful match (identity by state) between two alleles, and *n* is the total number of SNP sites in a pairwise comparison. The same equation was used to compute pIBS for an accession with its biological rep for error rate computation. In that case *i* and *j* represents the original accession and its biological replicate, respectively. Accessions with pIBS ≥99% (0.99) were grouped together in an arbitrary group number. Group size was computed as number of accessions in a group.

An error rate was computed using biological replicates for 76 accessions. Single to multiple seeds were grown for each accession, DNA was extracted, and sequencing performed as explained above. The error rate was computed using the following equation 2:

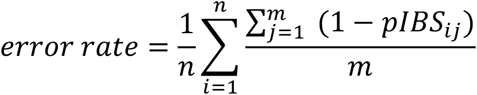

where, *n* is the number of accessions with biological replicates, *pIBS*_*ij*_ is the percent IBS for *i*th accession with its *j*th replicate, and *m* is the number of replicates for a given accession.

### Gliadin Profiling

To complement our GBS identity results, we extracted and profiled gliadin proteins from five independent groups of identical accessions (Supplementary Table S2) that were found to be the same with GBS. A single seed per accession was crushed in pestle and mortar to fine flour and mixed with 70% ethyl alcohol and stored at -4 °C for 24 hours. Following the protein extraction, samples were prepared using Bio-Rad Experion Pro260 kit (Bio-Rad, Hercules, California) following manufacturer’s instructions, and loaded on to an Experion Pro260 chip. The chips were read using Bio-Rad Experion automated electrophoresis system (Bio-Rad, Hercules, California). Virtual gel images were analyzed to compare accessions for identical protein banding patterns. For later comparison of protein profiling and GBS for two samples, multiple seeds were subjected to both procedures, where half of the seed was used for protein extraction and the other half with intact embryo was used for germination and tissue collection for DNA extraction.

### Imputing passport information

To facilitate the reduction of missing data and better curation of genebanks, we used genomic data and STRUCTURE^33^ software to impute the missing passport information for 26 WGRC accessions. For imputation, all the accessions with available passport information were used as learning samples and the remaining with missing to be imputed. The STRUCTURE parameters were set as follows: 10,000 burn-in iterations followed by 10,000 MCMC iterations, POPDATA=1, USEPOPINFO=1, GENSBACK=1, LOCIPOP=1, and all other parameters left at default settings. This resulted in posterior probabilities for each accession belonging to a specific geographical group with certain probability.

## RESULTS

### Sequencing and SNP genotyping

GBS generated ∼2 billion 100bp reads for Set 1, and DArTSeq generated ∼1 billion 77bp reads for Set 2, of which, 1.6 billion (83.4%) in Set 1 and 861 million (85.4%) contained expected sample barcodes followed by a restriction site. On average, each sample generated 1.9 million and 1.4 million barcoded reads for Set 1 and Set 2, respectively. Using these reads, discovery step in TASSEL-GBS pipeline found a total of ∼93 million unique 64bp tags. Each accession contributed an average of 81,365 unique tags that were aligned internally to find putative SNP sites, which resulted in 91,545 SNPs. Proportion of missing SNP data ranged from 0.6% to 78.9%. Population-level SNP filtering with ≤50% missing data, retained 29,555 SNPs that were used for cluster analysis. For pIBS, 20,844 pairwise comparisons were performed on average between any two accessions.

### Clustering and identifying identical accessions

Two different analyses were performed to identify identical accessions; a cluster analysis and allele matching. Cluster analysis (Supplementary Fig. S1) provides a quick method to cluster accessions based on genetic distances, however it cannot find identical accessions *per se*. For curating genebanks, cluster analysis should be used as a first step to group phenotypically cryptic accessions outside of the species under study and identify other outliers. From the cluster analysis, we observed the strong population structure between lineage 1 and lineage 2 that is known and previously reported in in *Ae. tauschii*^34^. As expected, we could assign all accessions into two large clusters, and identified three outliers which were removed from subsequent analysis (Supplementary Fig. S1). Accession TA3429 was found to be an outlier in STRUCTURE analysis. Two other accessions, one each from PAU and CIMMYT, clustered with TA3429 to form an outlier group. Corroborated by allele matching analysis, these outliers did not match with any other accession, supporting evidence that they have been either misidentified as *Aegilops tauschii* or could be the hybrids between two lineages.

Contrary to cluster analysis, allele matching provides an absolute percent IBS coefficient that can be used to identify identical accessions. Based on allele matching, different accessions had pairwise identity ranging from 37.5-99.9% (Supplementary Fig. S2). Each genebank resulted in a bimodal distribution of pIBS because of the strong population structure within *Ae. tauschii*. The higher pIBS peak represents the percent identity within subpopulations, and lower pIBS peak represents between subpopulations. With genotyping error, it is not possible to expect a 100% allelic identity for accession that should be considered the same. For this study, we implemented 99% allelic identity threshold for declaring accessions identical. This was initially based on expected sequencing error rates and confirmed with biological sample replicates. Minimum and maximum number of duplicated accessions were found in WGRC (25.88%) and PAU (54.01%), respectively, with CIMMYT having 43.04% duplicated accessions (Fig. 2). Combined across all genebanks, about 50% accessions were putatively duplicated. After removing the identical accessions, the WGRC, CIMMYT and PAU had only 421 (74.12%), 221 (45.99%) and 86 (45.99%) unique accessions, respectively. Based only on these unique accessions, pairwise IBS were computed for the accessions across the genebanks. The WGRC shared 32 (12.62%) with PAU and 129 (40.19%) accessions with CIMMYT, and PAU shared 29 (18.89%) accessions with CIMMYT. Overall, all three genebanks shared 26 (10.71%) accessions (Fig. 3) with group size of identical accessions ranging from 2 - 44 accessions (Supplementary Fig. S3). After grouping the accessions across all genebanks, only 564 unique accessions were found, representing over 50% duplicated accessions across the combined collections.

**Figure 2.**
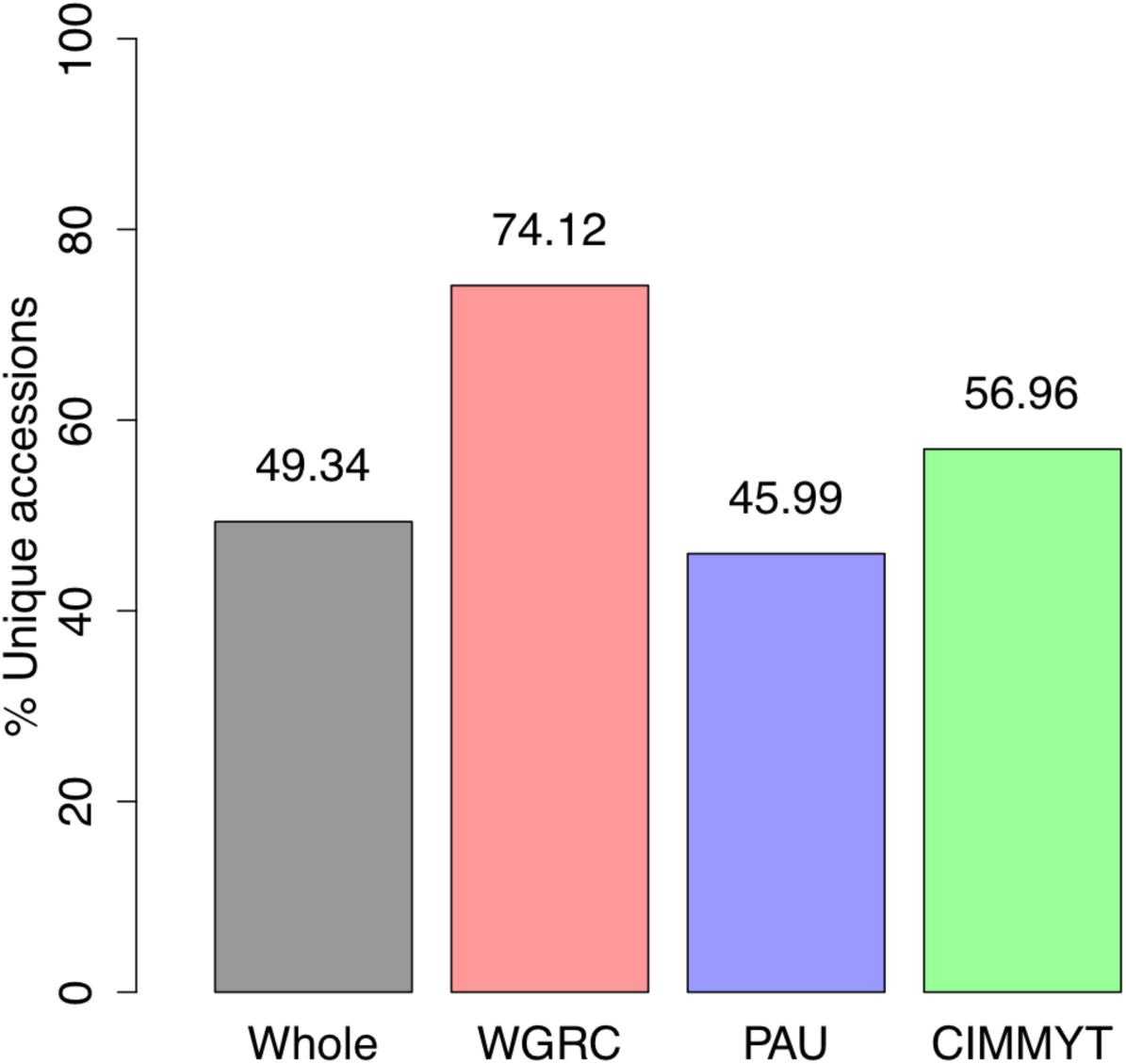
Bar plot showing percent unique accessions in whole collection, WGRC, PAU and CIMMYT genebanks. Values on top of each bar denotes the exact percent of unique accessions.

**Figure 3.**
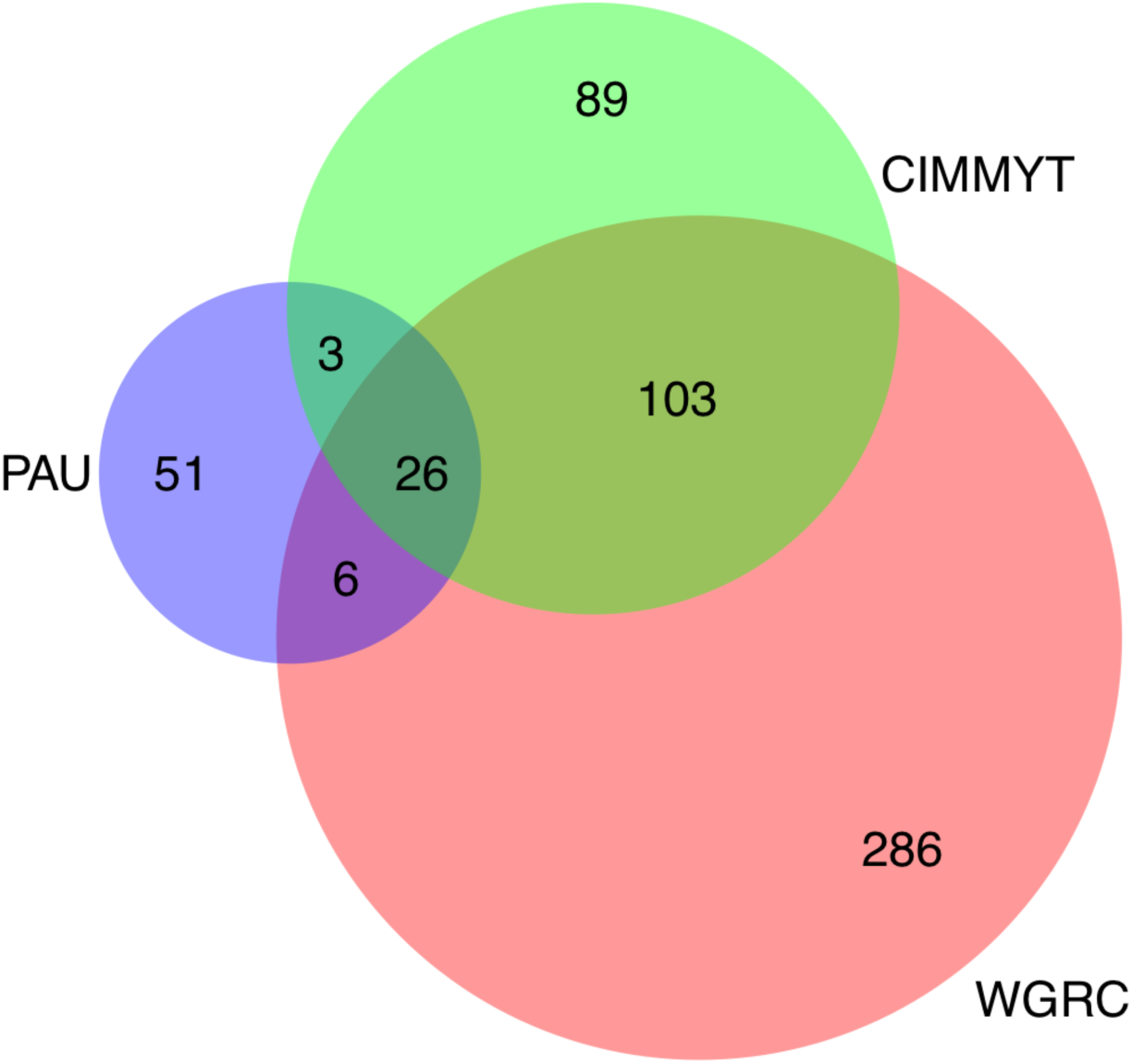
Venn diagram representing shared and unique accessions among and within genebanks. The total number for each genebank represents only unique accessions within a genebank.

### Error rate and efficiency

To compute the error rate of the GBS method, 76 accessions from the WGRC were resequenced and used as biological replicates. Of these 76 accessions, 11 had pIBS <99% with their respective original samples. Using the equation 2, the overall error rate was computed to be 3.13%, which is higher than our 1% threshold. To investigate further, multiple seeds from these 11 accessions were planted, however, only eight accessions produced at least one plant. GBS was performed on these eight accessions as described below.

Four out of eight accessions produced only a single plant. These were resequenced and compared with their previously sequenced respective samples (original sample and biological replicate). As initially expected, all four resequenced samples matched with >99% pIBS with either the original sample or the respective biological replicates. Two of these accessions matched with their original sample and other two matched with their biological replicates. These results point to the possibility of sample contamination that resulted in bad GBS data in one of the two initial GBS runs. Another possibility is that the original seed source was heterogeneous. Seed or sample mixture during the genotyping process of large number of samples is possible, however, we attempted to test the latter conjecture.

Remaining four out of eight accessions (TA1581, TA1589, TA1714 and TA2468) produced multiple plants that allowed us to test our hypothesis that the original seed source was heterogeneous. The final GBS was performed on each plant individually and compared with their respective original samples and biological replicates. TA1581 and TA1589 matched nicely with their original sample and all other replicates within this GBS run, but not the previous biological replicate. This points to the possibility that the sample contamination might have happened during the sequencing of previous biological replicates for these two accessions. In contrast, resequenced samples for TA2468 matched with >99% identity with the previous biological replicate and all other samples within this run, but failed to match with the original GBS. This again points to the possibility that the sample contamination might have happened during the original GBS.

For the final TA1714, a different pattern was observed. Two of the four resequenced samples matched with >99% identity with the original GBS, and the other two matched with the biological replicate. This supports our hypothesis and presents an evidence that the genebank seed source might be heterogeneous that results in lower pIBS. This is further evident in independent gliadin profiling discussed below. After removing these anomalous coefficients, the accuracy improved, and the error rate was reduced to only 0.48%, which is below our 1% threshold. This supports that the GBS is a robust tool for finding identical accessions in genebanks with very little error due to sequencing and biological sampling within homogenous accessions.

### Gliadin profiling

To independently validate the GBS results, gliadin profiling was run on eight independent groups from the cluster analysis in two separate runs. Gliadin proteins were selected for independent confirmation because of their ease of extraction and polymorphic profiling pattern. The first run included ten samples from four different groups (Supplementary Fig. S4). Per the manufacturer’s manual, bands lower than 10 kD were excluded as these are system bands that are produced by the small molecules interacting with lithium dodecyl sulfate (LDS) micelles in gel-staining solution and do not carry useful information. We observed matching banding pattern for the identical samples within the groups. For the second run (Supplementary Fig. S5), samples were included from four other different groups. As expected, the samples within all groups have similar banding pattern with the following notes. Sample TA2457 (Supplementary Fig. S5 - Lane 7) has the similar banding pattern as other samples from Grp15 (Lanes 5 and 6) but has a smeared profile that might be due to higher amount of extracted protein. Sample TA1579 (Supplementary Fig. S5, Lane 2) is the only accession from Grp187 and had very different banding pattern as compared to any other lane in this gel. Overall, matching banding pattern for the accessions within a group provides an independent evidence that the accession grouping based on GBS results are accurate.

### Detecting accession heterogeneity

TA1714 was hypothesized to be a heterogeneous and TA2457 a homogeneous accession based on the initial GBS grouping. To detect and confirm the heterogeneity in the source seed, these two accessions were subjected to a final GBS run. Half of the seed was crushed for protein extraction and the remaining half with intact embryo was germinated for tissue collection for GBS. For TA1714 and TA2457, 12 and 15 plants of each accession were planted, respectively, and subjected to GBS and gliadin profiling. As expected TA1714 showed heterogeneity in both the GBS and gliadin profiling by forming two sub-groups (Supplementary Fig. S6; red and blue box). Gliadin profiling was corroborated with GBS grouping of these samples. Contrary to TA1714, TA2457 did not show different banding pattern among individual plants from this accession (Supplementary Fig. S7), which supports that TA2457 is homogeneous. Both gliadin profiling results match with the corresponding GBS sub-groups. Independent confirmation with gliadin profiling supports that GBS can also be implemented to detect heterogeneity in the genebank samples.

### Imputing missing passport information

STRUCTURE analysis resulted in posterior probabilities ranging from 0.001-1. Higher posterior probability indicated higher likelihood that the accession belongs to a certain geographical group. Because these geographical groups are not completely isolated, we treated these groups as admixed populations, hence we used the posterior probability of 0.6 or more to assign an accession in a group. Using this analysis, we could assign 24 out of 26 accessions with missing geographical information into one of the geographic clusters. Two remaining accessions could not be assigned to any specific group because of lower probabilities (Supplementary Table S3).

## DISCUSSION

### Genotyping platform and accuracy

Selecting a genotyping platform is important when a large number of samples are of interest. We sequenced 1143 *Ae. tauschii* samples using two sequence-based methods. Sequence-based methods, such as GBS, are inexpensive and robust for genotyping a diverse range of uncharacterized species with complex genomes^12^. Here, we could use newly generated GBS data for set1 and previously generated DArTSeq for set2, to find duplicated accessions and efficiently curate the genebanks. As no prior SNP information is required for sequence-based methods, they also control for ascertainment bias because the SNP discovery and genotyping is performed on the same samples. Even though GBS only captured less than 1% of the genome, it resulted in an average of 20,844 pairwise SNP comparisons for allele matching. GBS grouping complemented with gliadin profiling, a very small error rate of only 0.48% makes it is a robust tool for this type of germplasm characterization.

### Collaborating with other genebanks

The ability to combine existing genotypic datasets and germplasm sharing is of great interest for genebank collaborations. As a starting point, this strategy was used on a diploid progenitor of wheat to identify unique accessions within and among genebanks. Here a coordinated effort between WGRC, CIMMYT and PAU could compare 1143 *Ae. tauschii* accessions across the genebanks and identify both identical and unique accessions across all the genebanks. Genebanks included in this study were rather smaller in size where all the accessions were genotyped and characterized, however, large scale genebanks usually lack this practice and record of duplicated accessions are often missing. Historically, even when these records are disseminated during germplasm sharing, they tend to lose track over time because of poor management practices. Therefore, the ultimate benefit of this strategy will be realized when this method is implemented globally in collaboration across all genebanks. The sequencing technology has quickly reached a point to enable globally coordinated effort among all genebanks to genetically curate these collections and find unique accessions in them. These globally unique accessions should then be prioritized and likely shared with other genebanks for additional backup of those irreplaceable accessions.

### Defining globally unique accessions

We have correlated many accessions with lost or incorrect accession identifiers through genotyping these collections. Most misclassifications happen during sharing of germplasm resources between collections^35^, which leads to significant duplication and incorrect information. Historically, germplasm was frequently shared, however, the associated metadata often was lost or misidentified, resulting in inaccurate classification and the new identifiers assigned lead to duplications in and across collections. Re-collecting at the same locations and sharing germplasm among genebanks also results in duplications within and among genebanks. We found 26-54% redundant accessions within, and a total of over 50% redundant accessions among the WGRC, CIMMYT and PAU genebanks. Our GBS results were corroborated by gliadin profiling. GBS generates genome-wide biallelic markers, whereas gliadin protein profiling samples multiple alleles from only a handful of loci, which complements and independently validating GBS results. As a starting point, we only performed this analysis for *Ae. tauschii*, but this strategy can be extended to other species with different ploidy levels stored in various genebanks. Genebanks worldwide are reported to hold over a million *Triticeae* accessions^36^. However, if our observations from this study hold true for other species, including the *Triticeae* tribe, we are vastly overestimating the number of unique germplasm accessions stored in the genebanks.

Applying genetic curation across genebanks around the world should be made a coordinated priority. Once unique accessions are identified across all collections, a globally unique ID could be generated and duplicate accessions within and between collections noted. With global curation, genebanks can better coordinate and curate collections efficiently. Currently, 482 genebanks use the GENESYS database (https://www.genesys-pgr.org) for over 3.6 million accessions which could provide a platform for establishing global curation. Such curation could also help other research endeavors, such as recently funded CGIAR Genebank Platform 2017-2022, whose main goal is to make available 750,000 accessions of crops and trees to the research community for crop improvement.

### Curating passport information and metadata

Often, vital metadata associated with shared germplasm, such as geographical or species information, is missing or incorrect. Species classification is a real challenge when dealing with cryptic species. A combination of existing genomic tools and statistical analyses can be used to infer those missing pieces. We used one such combination, GBS and cluster analysis and identified outliers (Supplementary Fig. S1). Although it is very difficult to accurately assign an accession to a geographical region at city level resolution, genotypic similarities and ancestry relationships can be used to group them together with other accession that have the metadata available. We used such methods to assign 24 out of 26 accessions to a potential geographical region of origin. Meyer (2015) noted that researchers tend to use germplasm with complete passport information and other associated metadata, which provides an incentive to collect and curate the accessions, and infer the missing information.

### Future direction for germplasm collection

The role of wild germplasm in crop improvement and the need to collect and preserve as much wild diversity as possible is evident. However, a specific protocol is necessary to avoid the accumulation of redundant accessions and keep only unique ones. One such approach is presented here (Supplementary Fig. S8). Briefly, when a new accession is collected or received, multiple seeds should be planted for tissue collection, and tissue should be collected in bulk from all plants, which was not ensured in this current study. We only sampled single seed from each accession, and it is possible that we missed within sample heterogeneity. Genotyping should be done on the bulked tissue from several seedlings. However, because *Ae. tauschii* is a highly self-pollinated species, it is very rare to find within accession heterogeneity unless due to seed mixture. Nevertheless, if possible, multiple independent samples should be sequenced for each accession. High level of heterozygous SNP calls, and mismatches within an accession, should point to the possibility of heterogeneous seed source that can be purified using single seed descent method. Bulked genotype data should be used for comparison to an existing genotyping database to find if the new accession is unique or identical to an existing collection. If unique, a new ID should be assigned, otherwise, the accession should be grouped together with the existing group of accessions. One such case study is explained below.

### Case study for collecting new accessions

About 92% of the WGRC accessions were collected in 1950s and 60s by various explorers and obtained through sharing among various genebanks. To fill the gaps in the collection sites and to preserve more genetic diversity, a recent collection expedition was conducted in June 2012 by WGRC researchers. During this expedition, a total of 44 accessions of *Ae. tauschii* were collected with passport information (blue dots; Fig. 1). Based on our analysis, only 36 collected accessions (∼66%) were unique in that they did not match with any other accession, either the newly collected or the already existing accessions. One newly collected accession had pIBS >99% with three already existing WGRC accessions that were collected decades ago. Seven accessions had pIBS >99% with at least one another new accession that were from the same geographic areas. Even though we collected 44 new accessions, but effectively only 36 (∼82%) of them were unique. These findings support implementation of a protocol for efficiently curating the genebanks in place, which is based on genotypic data.

## CONCLUSION

There are significant costs associated with running a genebank, beginning with acquiring an accession to storing and maintaining the germplasm. Because genebanks have limited funding and resources, identifying the duplicate accessions would result in a savings on both. Cost effective genotyping methods, such as GBS, can be applied for identifying duplicate accessions, and infer missing geographical and species information. Our results indicate that we are overestimating the diversity stored in the genebanks. Ultimately, identifying unique accessions within and across the genebanks will facilitate the better use of wild germplasm, make sharing more efficient, help breeders work with genetically diverse unique individuals and make better use of the untapped genetic diversity.

## Acknowledgements

This is contribution #18-400-J from the Kansas Agricultural Experiment Station, Manhattan. This study was conducted under the auspices of the Wheat Genetics Resource Center (WGRC) Industry/University Collaborative Research Center (I/UCRC) through support of industry partners and partially funded by NSF grant contract (IIP-1338897). We would also like to acknowledge the contribution of Industry Advisory Board to improve this manuscript with their valuable comments and suggestions.

## Author contributions

N.S. performed the analysis and wrote the manuscript; S.W. performed GBS and helped in other lab protocols; W.J.R. maintained and provided WGRC material; N.S., S.Se. and V.T. developed the idea; P.V. and S.Si. maintained and provided CIMMYT material; S.A. and P.C. maintained and provided PAU material; B.S.G. and J.P. conceived the idea, designed the experiment, directed the project and wrote the manuscript. All authors contributed to the review of this manuscript.

## Data availability

Sequence reads generated using genotyping-by-sequencing are available from NCBI SRA under accession SRP141206 and R-code is available from the corresponding author upon request.

---

**Figure S1.**
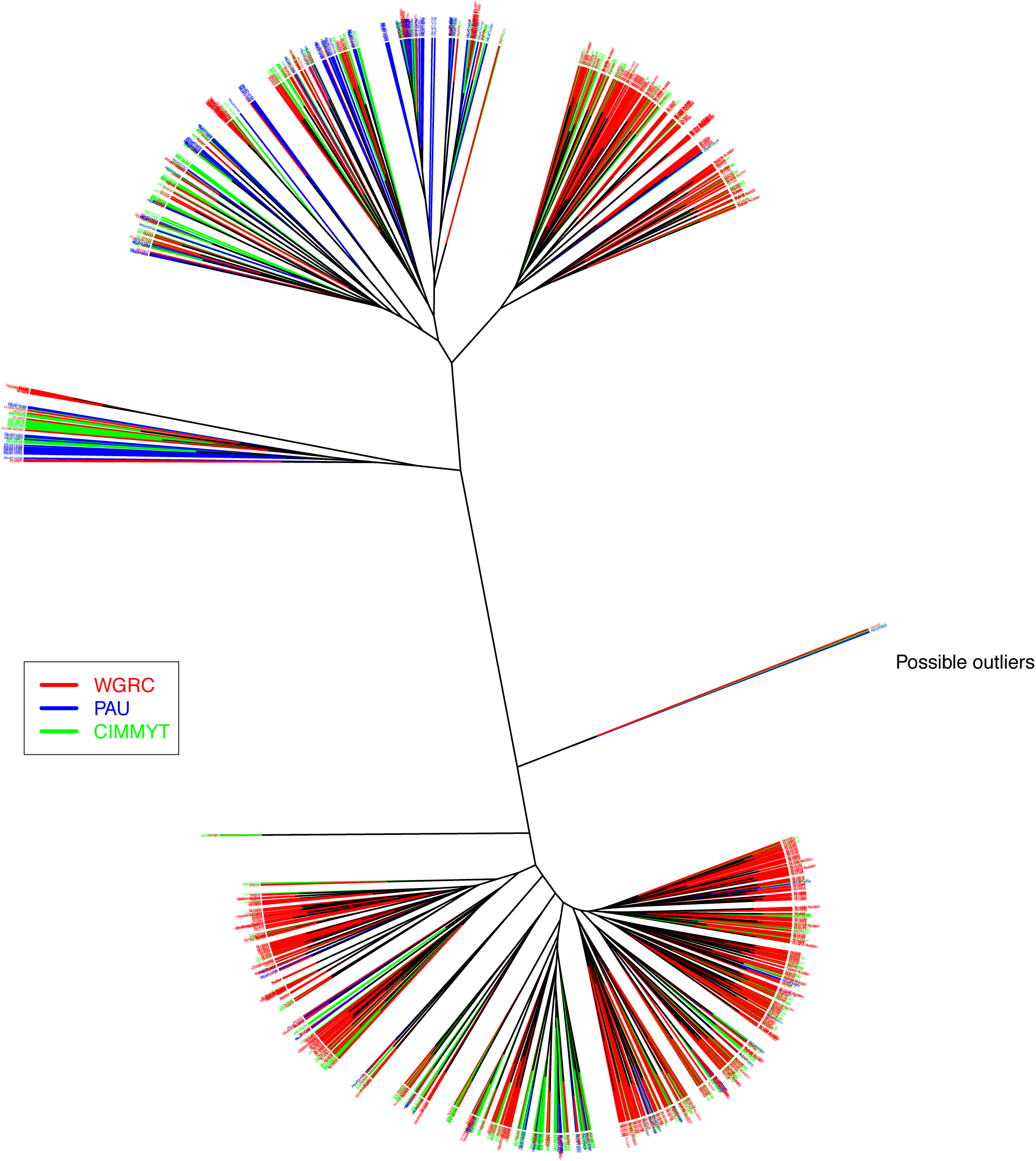
Cluster analysis of WGRC, PAU and CIMMYT genebanks’ *Aegilops tauschii* collection using genotyping-by-sequencing.

**Figure S2.**
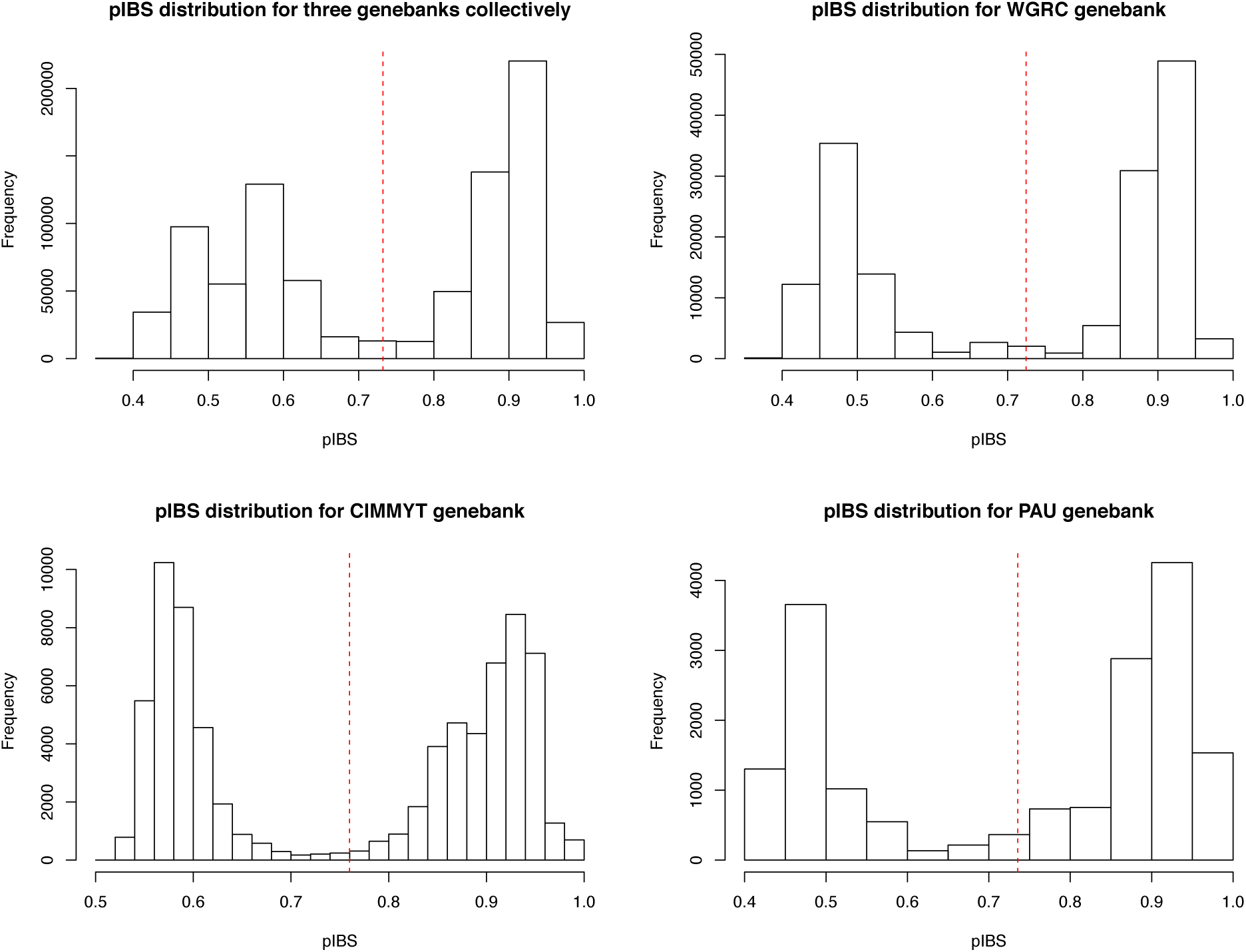
Percent identity by state (pIBS) distributions for three genebanks separately and collectively.

**Figure S3.**
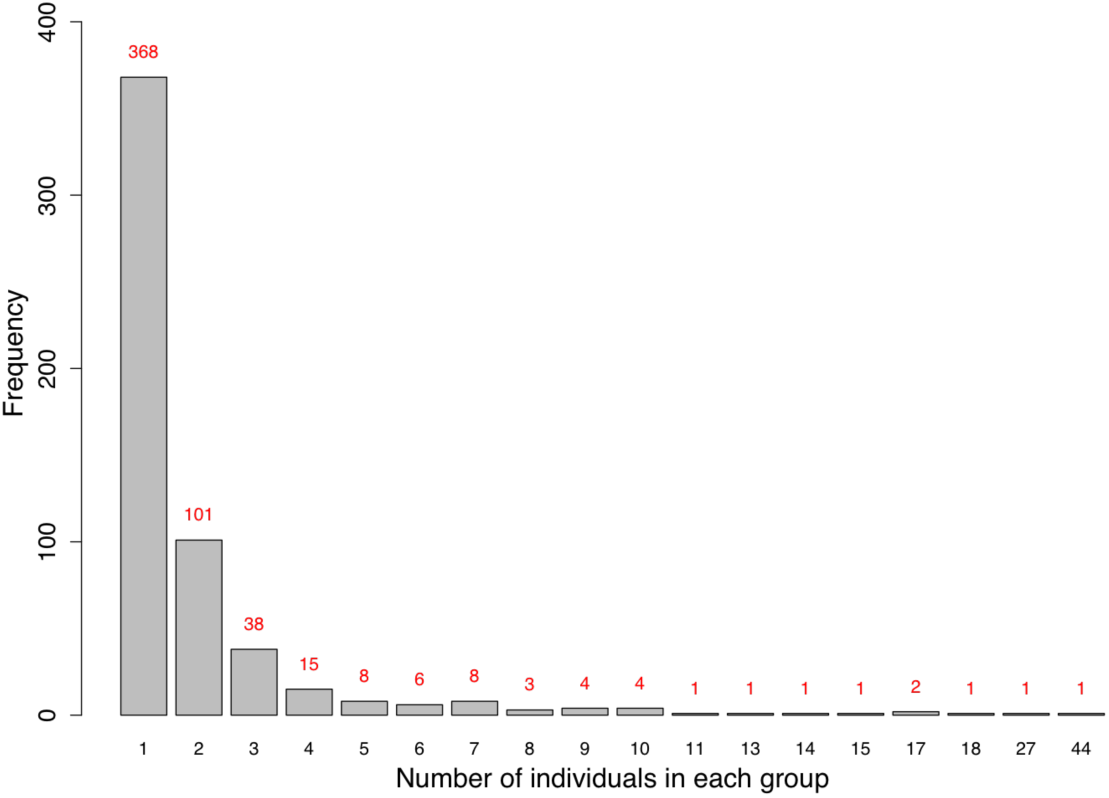
Bar plot showing frequency of each group size. Values on top of each bar represents the exact frequency of corresponding group size listed on x-axis. Total of 368 accessions were total unique and did not match with any other accession.

**Figure S4.**
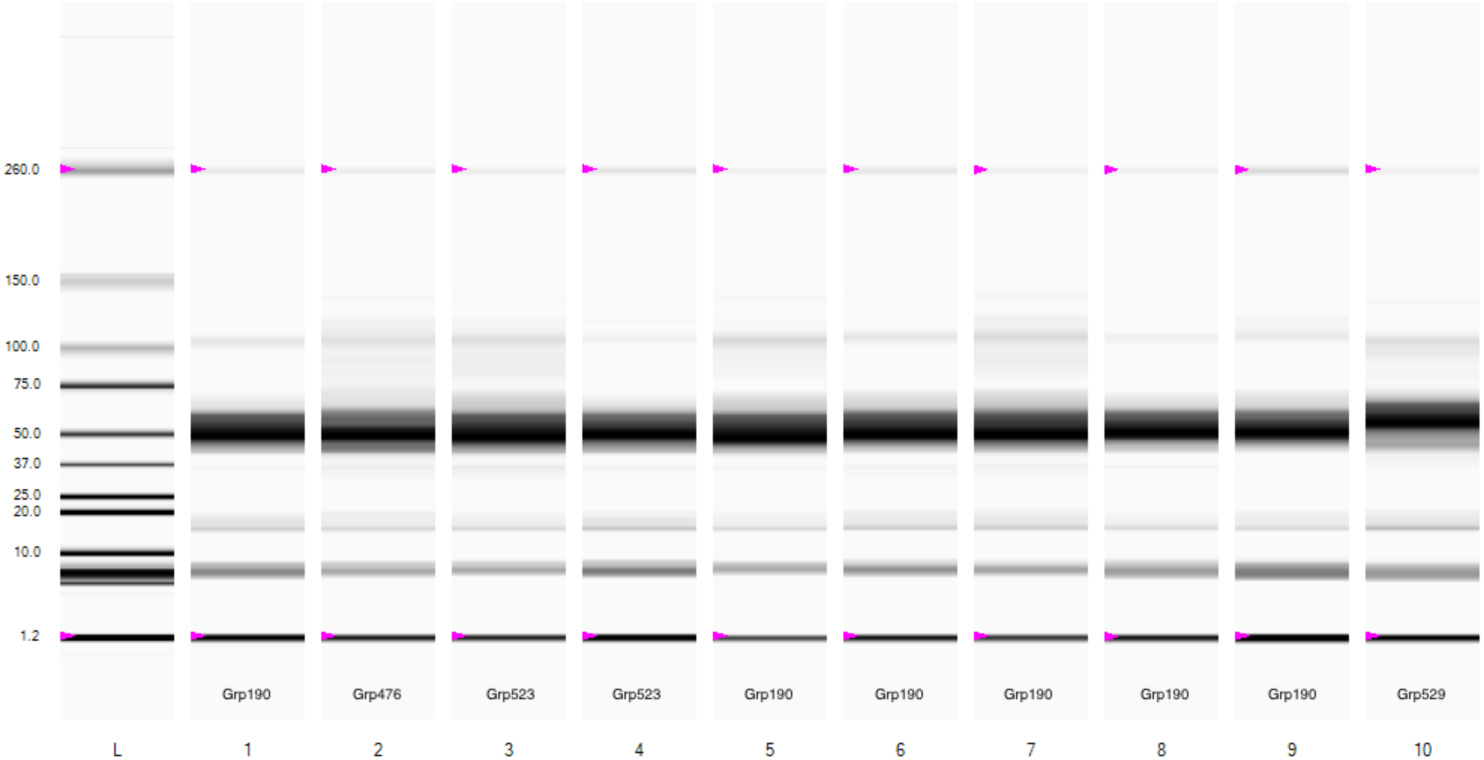
Virtual gel image showing accessions from four different groups (lanes 1 and 5-9 from Grp190, lane 2 from Grp476, lanes 3-4 from Grp523 and lane 10 from Grp529). As expected, lanes 2 and 10 shows different banding pattern as they are the only representative of their respective groups on this gel. Lanes 3 and 4 have similar banding pattern. Lanes 1 and 5-9 from Grp190 have similar banding pattern. This suggests that accessions within a group tend to have a similar banding pattern, which corroborates with the accession grouping with allele matching.

**Figure S5.**
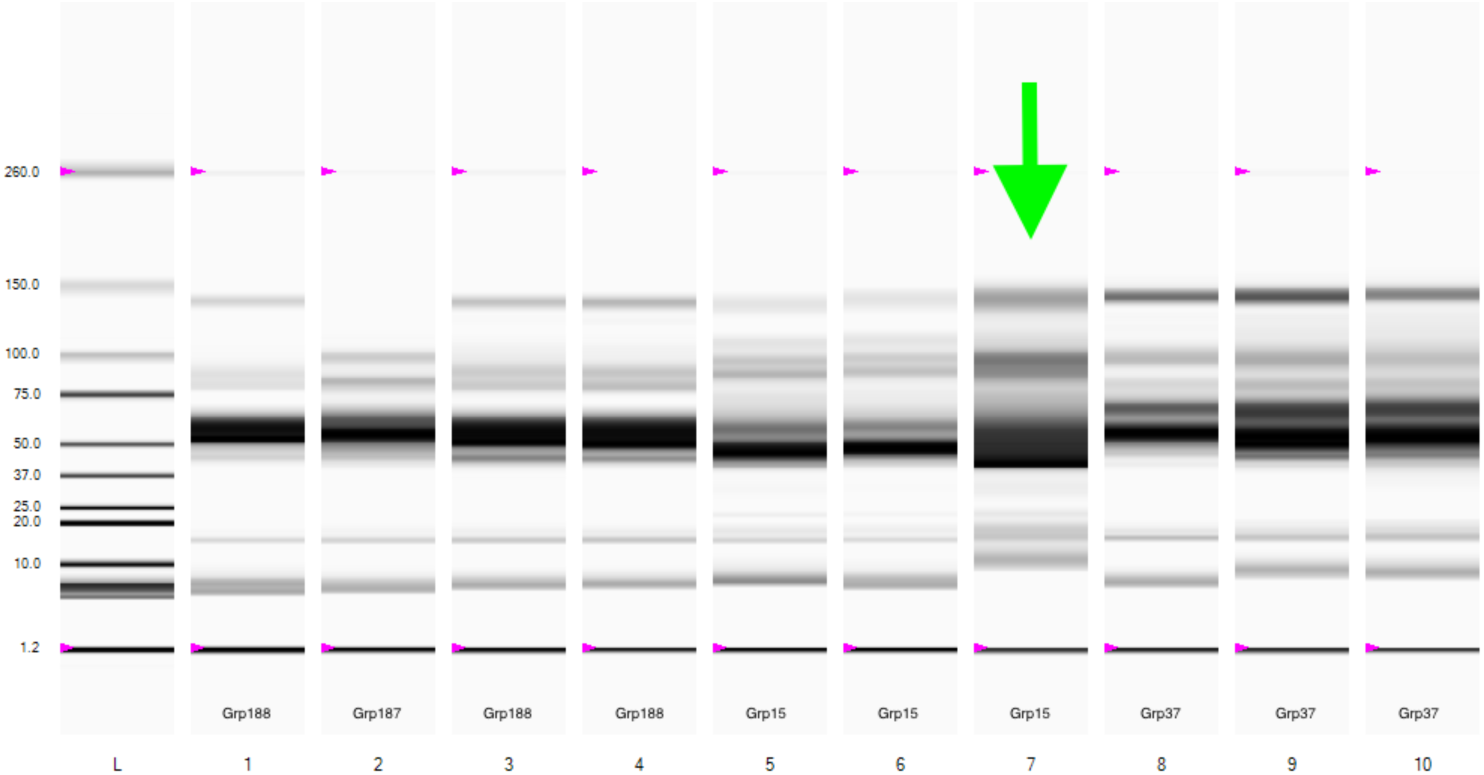
Virtual gel image showing accessions from four different groups (lanes 1, 3, and 4 are from Grp188; lane 2 from Grp187; lanes 5-7 from Grp15; and lanes 8-10 from Grp37). Lanes 1,3 and 4 have similar banding pattern; lane 2 has totally different banding pattern not matching with any other lane; lanes 5-7 have similar banding pattern but lane 7 (green arrow) seems to have very high concentration of the protein, giving it a smear look; lanes 8-10 seem to have similar banding pattern. This suggests that accessions within a group tend to have a similar banding pattern, which corroborates with the accession grouping with allele matching.

**Figure S6.**
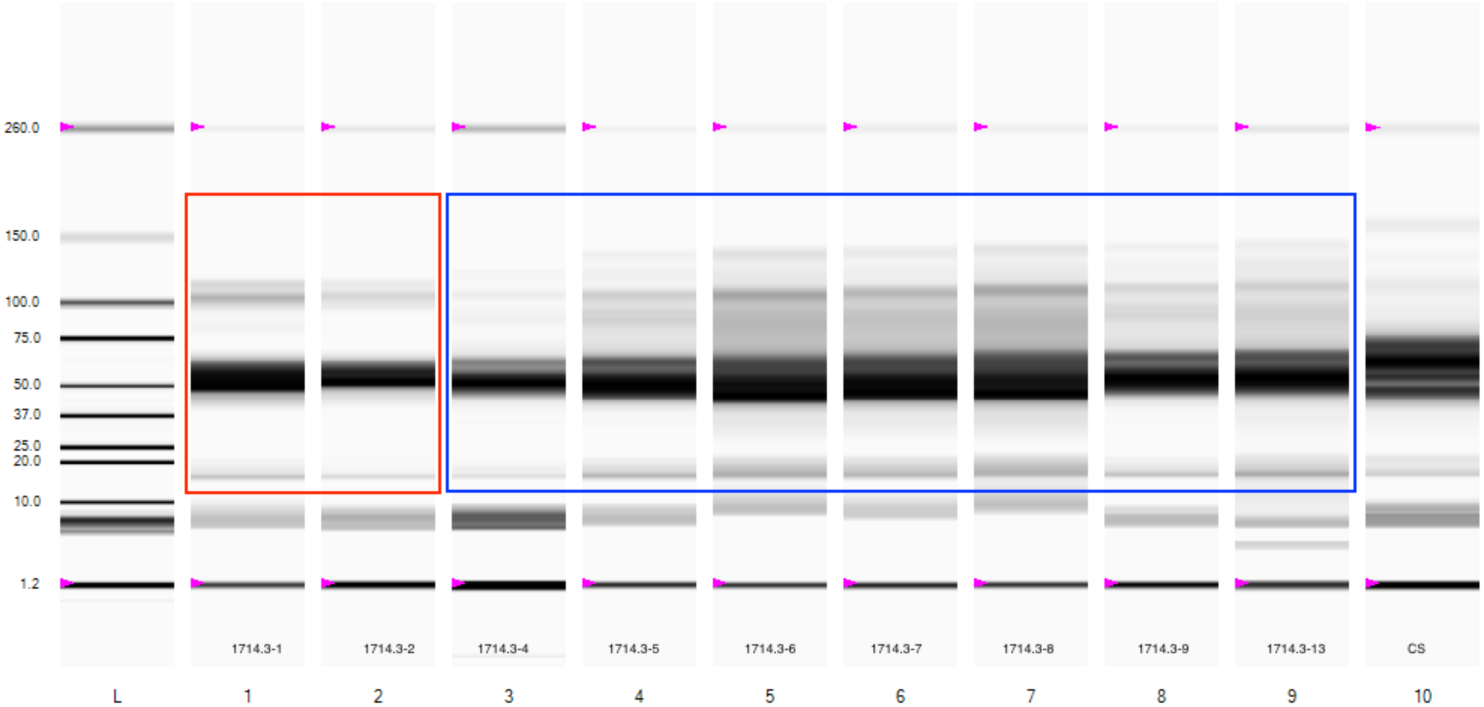
Virtual gel image showing gliadin profiling for heterogeneous accession TA1714. First two lanes (red box) have a similar banding pattern forming a group, and lanes 3-9 (blue box) have similar banding pattern with minor differences. Lane 10 is Chinese spring wheat for control. The different patterns between red and blue box samples presents an evidence that the samples came from a heterogeneous seed source.

**Figure S7.**
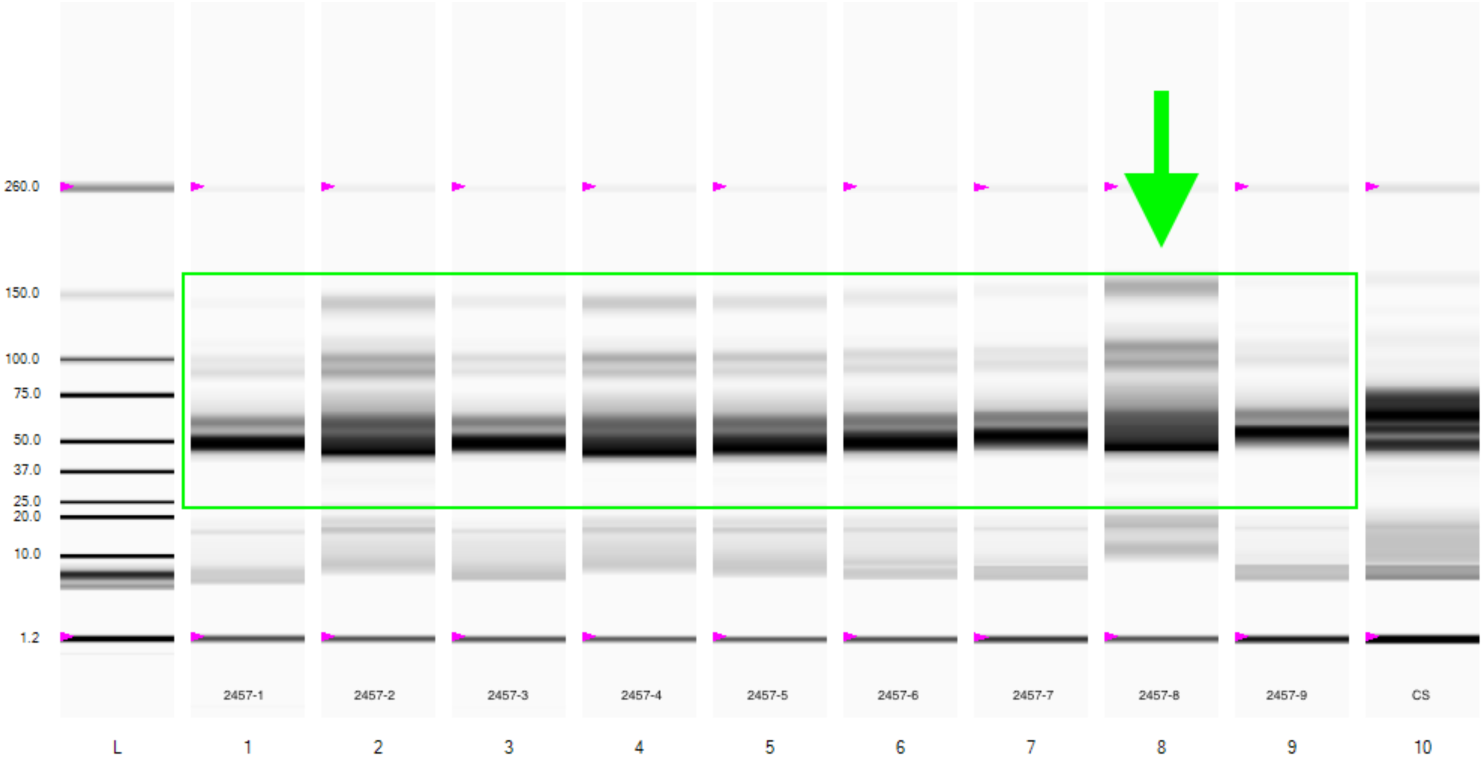
Virtual gel image showing gliadin profiling for homogeneous accessions TA2457. With some minor differences, banding pattern for lanes 1-9 (green box) look similar with an exception of lane 8 (green arrow). Sample in lane 8 does appear to have a similar banding pattern but possibly has higher extracted protein concentration that gives it a smeared look.

**Figure S8.**
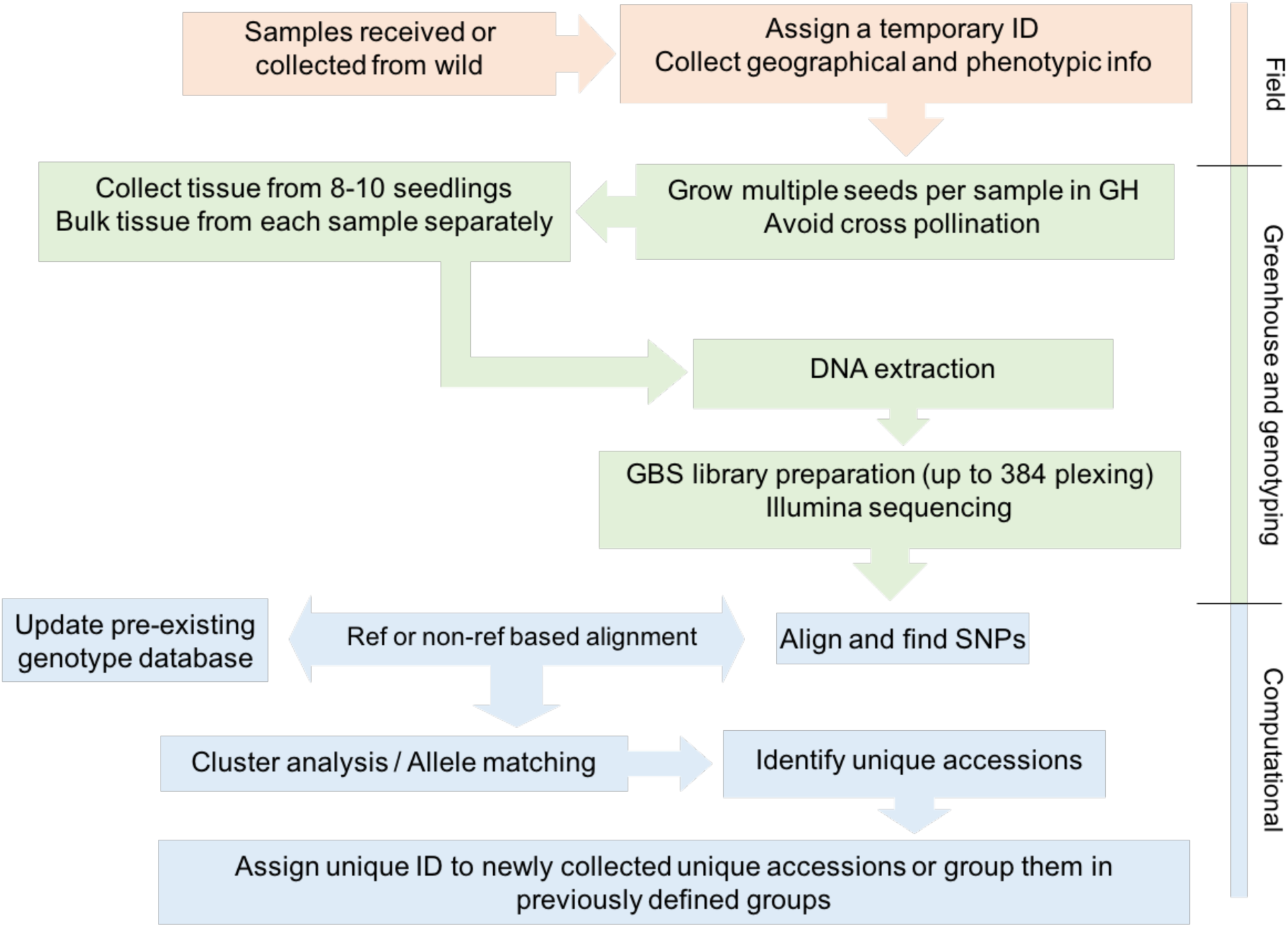
Future germplasm collection and management strategy to avoid the accumulation of redundant germplasm accessions.

